# Stable Genetic Transformation and Heterologous Expression in the Nitrogen-fixing Plant Endosymbiont *Frankia alni* ACN14a

**DOI:** 10.1101/496703

**Authors:** Isaac Gifford, Summer Vance, Giang Nguyen, Alison M Berry

## Abstract

Genus *Frankia* is comprised primarily of nitrogen-fixing actinobacteria that form root nodule symbioses with a group of hosts known as the actinorhizal plants. These plants are evolutionarily closely related to the legumes, which are nodulated by the rhizobia. Both host groups utilize homologs of nodulation genes for root-nodule symbiosis, derived from common plant ancestors. However the corresponding endosymbionts, *Frankia* and the rhizobia, are distantly related groups of bacteria, leading to questions of their symbiotic mechanisms and evolutionary history. To date, a stable system of genetic transformation has been lacking in *Frankia*. Here, we report the successful electrotransformation of *Frankia alni* ACN14a, by means of replicating plasmids expressing chloramphenicol-resistance for selection, and the use of GFP as a marker of gene expression. We have identified type IV methyl-directed restriction systems, highly-expressed in a range of actinobacteria, as a likely barrier to *Frankia* transformation and circumvented this barrier by using unmethylated plasmids, which allowed the transformation of *F. alni* as well as the maintenance of the plasmid. During nitrogen limitation, *Frankia* differentiates into two cell types: the vegetative hyphae and nitrogen-fixing vesicles. When the plasmid transformation system was used with expression of *egfp* under the control of the *nif* gene cluster promoter, it was possible to demonstrate by fluorescence imaging the expression of nitrogen fixation in vesicles but not hyphae in nitrogen-limited culture.

**Importance:** To date, the study of *Frankia*-actinorhizal symbioses has been complicated by the lack of genetic tools for manipulation of *Frankia*, especially stable genetic transformation. The transformation system reported here, particularly coupled with marker genes, can be used to differentiate patterns of gene expression between *Frankia* hyphae and vesicles in symbiosis or in free-living conditions. This will enable deeper comparisons between *Frankia*-actinorhizal symbioses and rhizobia-legume symbioses in terms of molecular signaling and metabolic exchange that will broaden understanding of the evolution of these symbioses and potentially make possible their application in agriculture. The development of transformation methods will allow further down-stream applications including gene knock-outs and complementation that will, in turn, open up a much broader range of experiments into *Frankia* and its symbioses.

## Introduction

Bacteria in the genus *Frankia* form nitrogen-fixing root nodule symbioses with a group of host plants, the actinorhizal plants. The actinorhizal plants are evolutionarily closely related to the legumes within the Nitrogen-fixing Clade (NFC; Soltis *et al*., 1995). The bacterial symbionts are only distantly related however: *Frankia* belongs to the phylum Actinobacteria, comprised of high-GC-content gram-positive bacteria, whereas the rhizobia, symbionts of the legumes, are gram-negative proteobacteria. Additionally, *Frankia* is a multicellular, hyphal genus whereas the rhizobia are single-celled. In most of the actinorhizal symbioses, and in nitrogen-limiting conditions *in vitro*, *Frankia* differentiates into two cell types: vegetative hyphae and nitrogen-fixing vesicles. Vesicles are surrounded by a lamellar hopanoid lipid envelope (Berry *et al.,* 1993) that increases in number of layers in response to oxygen tension thereby likely reducing the flow of oxygen into the vesicle interior to protect the nitrogenase complex from deactivation (Benson and Silvester, 1993). Thus, unlike the rhizobia which require their hosts to regulate the flow of oxygen for nitrogen fixation in symbiosis (Appleby, 1984), *Frankia* vesicles are capable of fixing nitrogen in atmospheric oxygen conditions (Benson and Silvester, 1993).

Biological nitrogen fixation has potential applications in the development of minimal input strategies for more sustainable agriculture globally and in nutrient-deficient soils through genetic improvement of crops (Gutierrez, 2012). In efforts to understand the mechanisms of root nodule symbioses, phylogenetic studies of the Nitrogen-Fixing Clade have shown a shared predisposition underlying the evolution root nodule symbioses between the actinorhizal plants and legumes (Werner *et al*., 2014; Battenberg *et al*., 2018). The actinorhizal and legume hosts share the Common Symbiotic Pathway, a signaling pathway that leads to the development of the nodule in response to the symbiont (reviewed in Oldroyd 2013) as well as several key gene orthologs in the process of nodule development (Battenberg *et al.,* 2018; Griesmann *et al*., 2018). Clear differences also exist in the genetics of nodulation in the two bacterial groups, leaving major questions concerning the evolution and development of root-nodule symbioses in the bacterial partners. The rhizobial signaling molecules that trigger nodulation in many of the legume hosts, called Nod factors, are synthesized by *nod* genes and have been extensively studied (Oldroyd, 2013). The majority of *Frankia* genomes do not contain clear sets of *nod* gene homologs with the exception of several of the cluster II *Frankia*: genomes of these *Frankia* contain homologs of the rhizobial *nodABC* genes that are expressed in symbiosis (Persson *et al*., 2015). Other mechanisms have been found to trigger legume nodulation by rhizobia, including an effector protein injected into the host by a type III secretion system (Okazaki *et al*., 2013); however genes responsible for these pathways lack homologs in *Frankia* genomes as well. Signals from *Frankia* in clusters I and III that elicit host root responses have been detected experimentally, but the chemical composition has not been defined (Ceremonie *et al*., 1999; Cissoko *et al*., 2018).

The major advances in knowledge of rhizobial symbioses have been made possible by the development of genetic tools for dissecting the metabolic and regulatory pathways in the microsymbiont-host interactions. Long *et al*. (1982) originally demonstrated the role of the *nod* genes in symbiosis by complementing nodulation-deficient mutants of *Sinorhizobium meliloti* with *nod* genes encoded on a replicating, broad host-range, plasmid. More recently, expression of marker genes in rhizobia has enabled a wider-range of experiments including the tracking of microsymbionts during the host infection process (Gage, 2002) and the identification of regulatory networks involved in symbiotic interactions (Spaepen *et al*., 2009). However, to date, there has been no stable genetic transformation system for *Frankia* (Kucho *et al*., 2009). As reported here, a transformation system in *Frankia* will thus enable functional inquiries into the diverse mechanisms involved in establishing root-nodule symbiosis and maintaining biological nitrogen fixation in actinorhizal symbioses. These inquiries will, in turn, contribute to a wider understanding of the origins of root nodule symbioses and mechanisms of *Frankia* and rhizobia interaction with hosts in the NFC.

*Frankia* has several barriers to transformation. Actinobacteria in general have low rates of homologous recombination due to competition between their homologous recombination pathway and a Non-Homologous End-Joining pathway (Zhang *et al*., 2012). Additionally, natural vectors for *Frankia* transformation are limited as no *Frankia* phages have been discovered (Simonet *et al*., 1990). Finally, its multi-cellular hyphae require that all cells be transformed in order for experiments to be viable. Despite these barriers, it has been demonstrated that DNA can be electroporated into *Frankia* cells (Myers and Tisa, 2004), providing a potential avenue for genetic transformation. Furthermore, Kucho *et al.* (2009) reported initial success in integrating a non-replicating plasmid into the chromosome of *Frankia* sp. CcI3. However the recombined plasmid was lost in the following generation, limiting its use in experiments.

Restriction enzymes pose a barrier to successful transformation in bacteria due to their role in defense by digesting foreign DNA. In some actinobacteria, the use of unmethylated plasmids has increased transformation efficiency (Ankri *et al*., 1996; Molle *et al*., 1999), potentially due to the presence of Type IV methyl-directed restriction enzymes. In the majority of bacterial taxa, DNA is methylated during replication by the methyltransferase Dam to mark parent strands for DNA repair and excision of misincorporated bases from the daughter strand (Sanchez-Romero *et al*., 2015). The majority of actinobacteria, however, lack *dam* homologs (Sanchez-Romero *et al*., 2015) and an investigation of genomes of *Streptomyces*, *Rhodococcus*, and *Micromonospora* spp. found that these genomes were not methylated in the canonical Dam pattern (Novella *et al*., 1996). While actinobacteria have recently been shown to have a unique mismatch repair pathway (Castaneda-Garcia *et al*., 2017) it is not yet known how this pathway utilizes methylation, if at all.

In this study we report successful genetic transformation of *Frankia alni* ACN14a by using an unmethylated plasmid to circumvent methylation-targeting restriction enzymes encoded in the ACN14a genome. This permitted the maintenance of plasmids derived from one with a very broad host-range origin of replication (Kurenbach *et al*., 2003) in *F. alni* cultures. Additionally, we show that *gfp* expressed from a replicating plasmid can be used to label gene expression differences between *Frankia* cell types during nitrogen fixation, opening up molecular genetics-based experiments on *Frankia* symbioses in the future.

## Results

### Identification of Restriction Enzymes in Frankia and Transcriptome Analysis

Type I, II and IV restriction enzymes were identified in several published *Frankia* genomes (Figure 1). Examining the transcriptome of *Frankia alni* ACN14a in (+)N culture (Alloisio *et al*., 2010), we found that three restriction enzyme genes were highly expressed: one type I enzyme (FRAAL4992, 92^nd^ percentile), one type II enzyme (FRAAL0249, 91^st^ percentile) and one type IV enzyme (FRAAL3325, 85^th^ percentile) (Figure 1). The type IV enzyme was annotated in REBASE as a “Type IV Methyl-directed restriction enzyme” of the Mrr methyladenine-targeting family. Other *Frankia* genomes contained likely homologs of this type IV enzyme as well. The genome of *Frankia casuarinae* CcI3 contained five type IV enzymes, the most of any *Frankia* genome examined (Figure 1). In the *Frankia* sp. CcI3 transcriptome, type IV Mrr enzymes were very highly expressed, up to the 96^th^ percentile. One of the five enzymes in CcI3 (Francci3_2839) was annotated as a Mcr type IV enzyme, which targets 5-methylcytosine instead (REBASE) and its expression was very low, in only the 18^th^ percentile of genes in the transcriptome. Of the complete genomes examined, only *Candidatus* Frankia datiscae Dg1 did not contain any putative type IV restriction genes.

**Figure 1.**
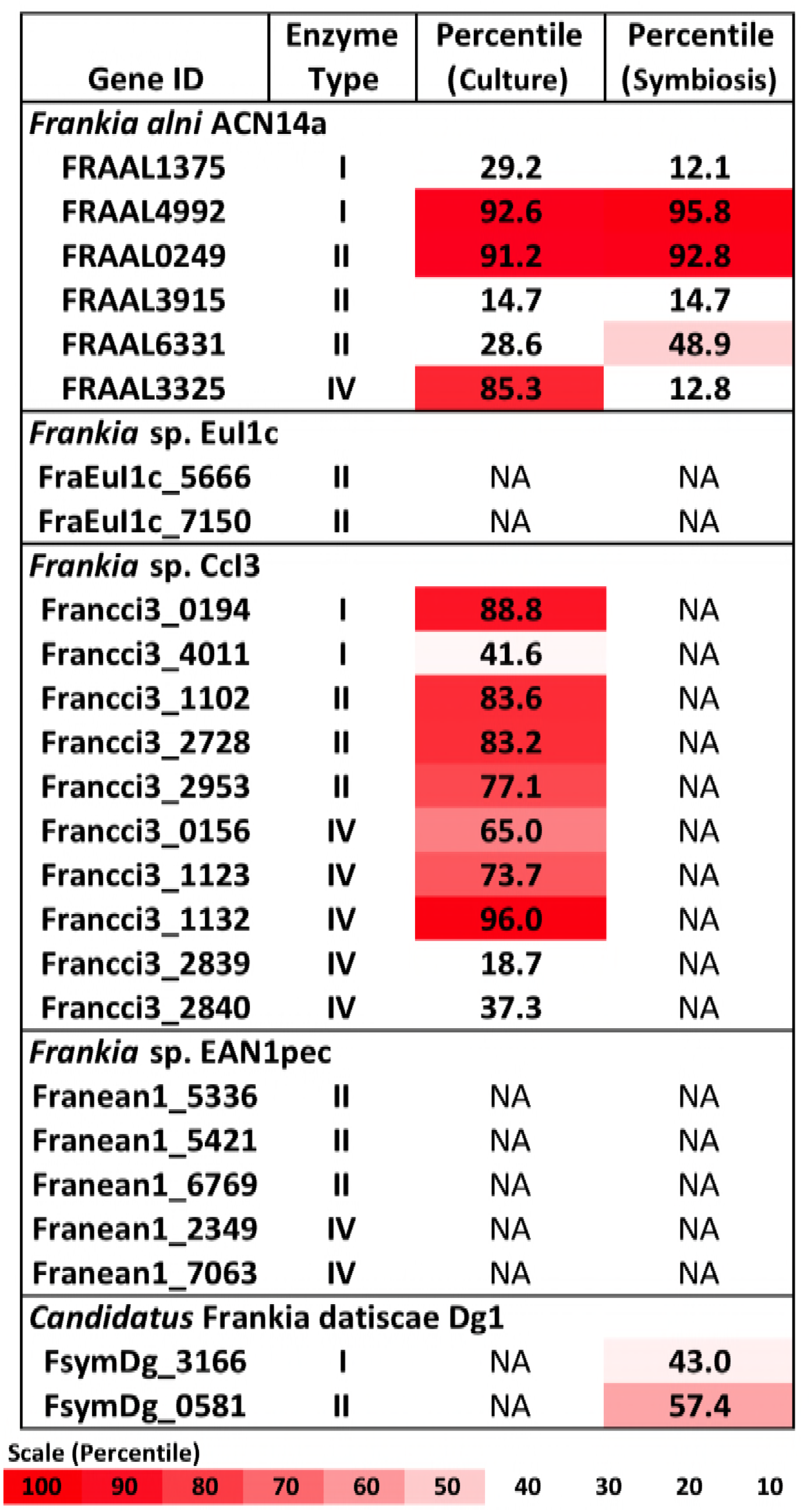
Restriction enzymes annotated in the *Frankia* genomes and expression levels of each gene in the transcriptome in culture when available (Alloisio *et al*., 2010 and Bickhart and Benson, 2011). NA: Transcriptome not available.

A symbiotic transcriptome of *F. alni* ACN14a from root nodules is available in addition to transcriptomes of free-living culture (Alloisio *et al*., 2010). In symbiosis with *Alnus glutinosa* the *mrr* homolog was down-regulated relative to the rest of the transcriptome, from the 85^th^ percentile in culture to an expression level at maximum no higher than 12 percent of transcriptome genes, while the other restriction enzymes identified in the *F. alni* genome were not down-regulated in symbiosis (Figure 1). In the other *Frankia sp*. with a symbiotic transcriptome available, the uncultured cluster II species *Frankia* sp. Dg1 (Persson *et al*., 2015), expression of its two restriction enzymes, neither of which was a Type IV enzyme, occurred at very low levels (Figure 1).

Of the actinobacteria outside of genus *Frankia* with published transcriptomes in pure culture, *Mycobacterium smegmatis*, *Streptomyces avermitilis*, and *Rhodococcus jostii* also highly expressed type IV restriction enzyme genes, in the upper-70^th^ percentiles of their respective transcriptomes (Figure 2). In comparison, the transcriptomes of proteobacteria and firmicutes examined expressed their corresponding *mrr* genes around the 50^th^ or 60^th^ percentiles. One notable exception was the actinobacterium *Mycobacterium tuberculosis*, which expressed its single gene for a methyladenine-directed restriction enzyme (Mrr) at extremely low levels, around the 6^th^ percentile in culture.

**Figure 2.**
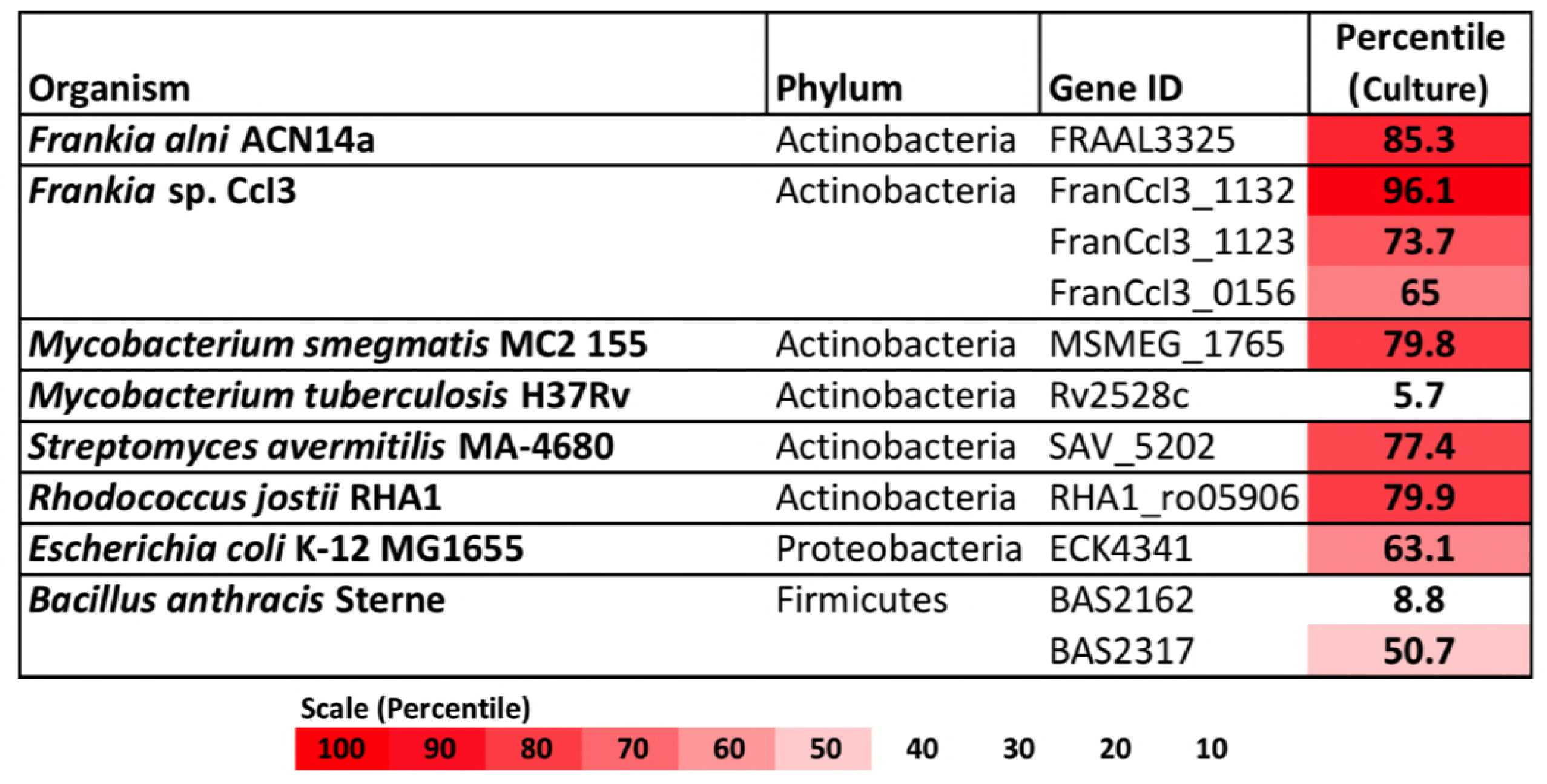
Restriction enzymes annotated in the genomes of *Frankia*, other actinobacteria, and proteobacteria and firmicutes for comparison. For each, expression levels for available transcriptomes growing in pure culture are provided.

### Genetic Transformation of Frankia alni ACN14a with an Unmethylated Replicating Plasmid

After electroporation *F. alni* cells formed visible hyphae in culture after approximately ten days (data not shown). When sub-cultured into chloramphenicol-selective media *F. alni* cultures transformed with unmethylated plasmid pSA3 were able to grow, whereas those transformed with methylated plasmid were not. When imaged under 488nm wavelength of excitation in the confocal microscope, green fluorescence typical of GFP was observed in hyphae as shown in Figure 3. Wild-type *F. alni* hyphae displayed autofluorescence around 575nm (data not shown), as observed by Hahn *et al.* (1993), but no fluorescence was observed in the 500-550nm range (Figure 3).

**Figure 3.**
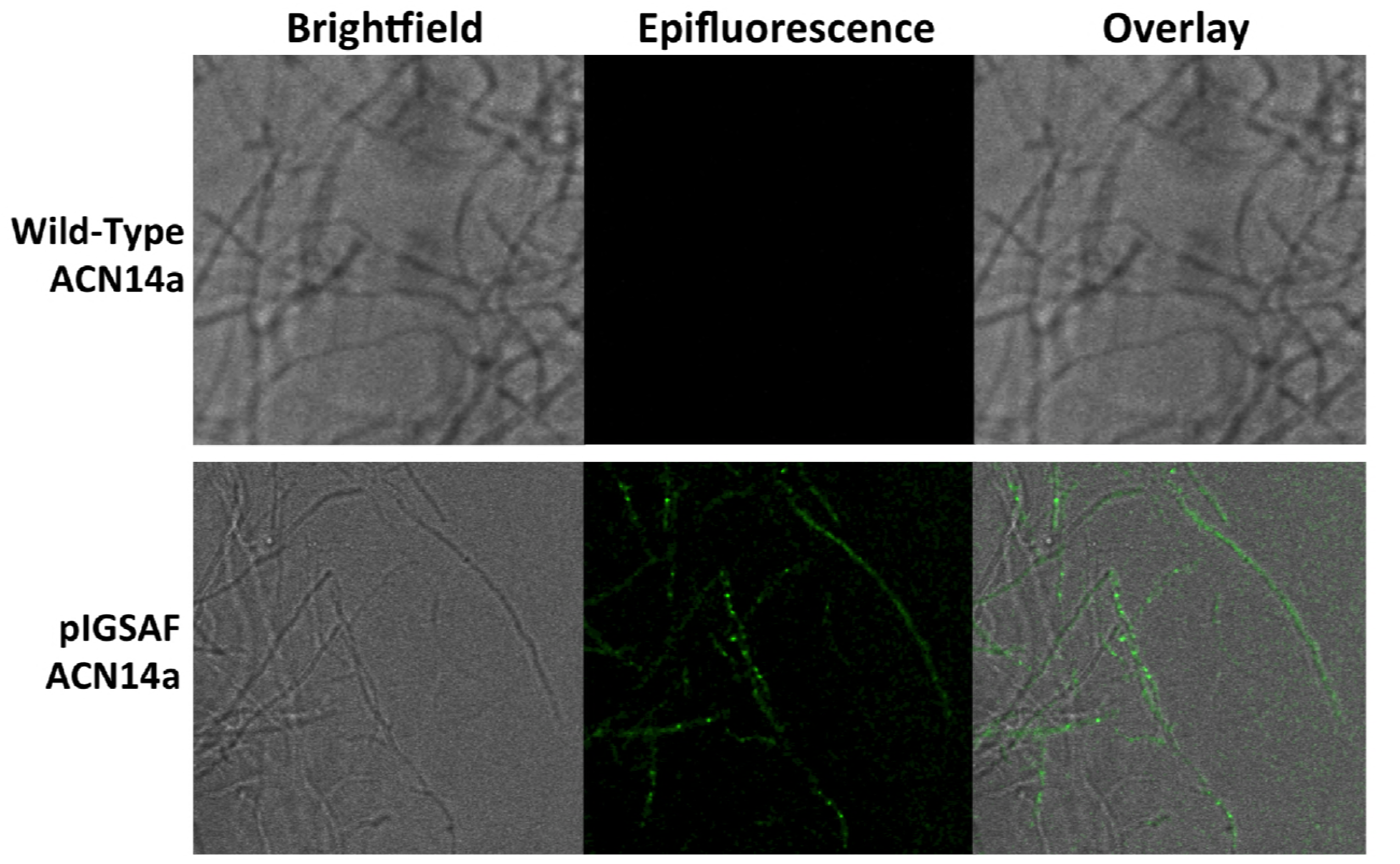
Transformation of *Frankia alni* ACN14a with a plasmid expressing *egfp*. Top: Wild-type *F. alni* ACN14a control. Bottom: Transformed *F. alni* ACN14a with plasmid pIGSAF. Images were obtained on a confocal microscope with both brightfield and epifluorescence and then overlaid.

### Differential Regulation of egfp Under the Control of the Frankia alni ACN14a nif Cluster Promoter

When grown in (–)N culture, transformants carrying the pIGSAFnif plasmid showed significant up-regulation of the *egfp* gene conjugated to the *nif* cluster promoter, approximately 100-fold relative to (+)N culture (Table 1). The *nifH* gene, used as a positive control for nitrogen fixation, was also significantly up-regulated approximately 9-fold in (–)N media compared with (+)N cultures. Thus, the up-regulation of the egfp gene is consistent with the previous report that plasmid pSA3 is maintained at approximately 10 copies per cell (Dao and Ferretti, 1985). Expression of the *rpoD* housekeeping gene, used as a negative control, was not significantly different between (+)N and (–)N cultures.

**Table 1.** qPCR verification of differential regulation of *egfp* in (+)N and (–)N media. qPCR tested fold-change of *egfp* as well as *nifH* as a nitrogen-fixation positive control and *rpoD* as a housekeeping negative control.

| Gene | Role | Fold Change<br>(-N/+N) | p-value |
| --- | --- | --- | --- |
| *nif:egfp* | Fluorescence | 101.6 | <.05 |
| *nifH* | Nitrogen fixation | 8.5 | <.05 |
| *rpoD* | Housekeeping | 0.4 | >.05 |

*F. alni* containing pIGSAFnif grown in (–)N media fluoresced predominantly in the vesicles (Figure 4). Little to no fluorescence was observed in hyphae. Fluorescence in the vesicles was present both in the spherical portion as well as in the stalk connecting the vesicle to the hyphae. No fluorescence was observed in hyphae or vesicles of wild-type *F. alni* grown in (–)N media.

**Figure 4.**
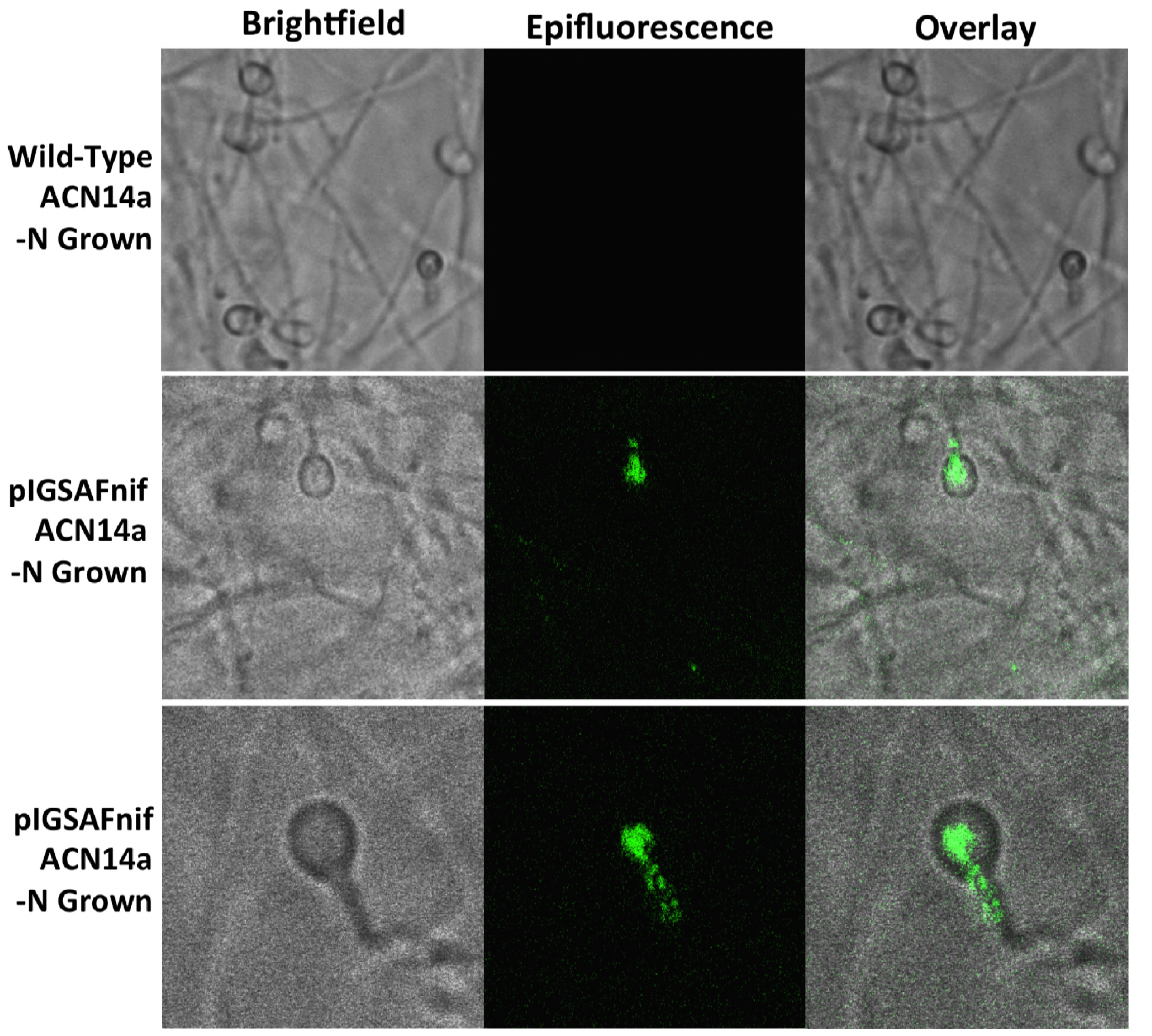
Transformation of *Frankia alni* ACN14a with a plasmid expressing *egfp* under the control of the *nif* cluster promoter region from the *F. alni* genome. Transformed cultures were grown in both (+)N and (–)N media to compare fluorescence with and without nitrogen fixation. Top: *F. alni* ACN14a transformed with plasmid pIGSAFnif grown in (+)N media. Middle: *F. alni* ACN14a with plasmid pIGSAFnif grown in (–)N media. Bottom: Close-up of a vesicle showing fluorescence in the stalk. Images were obtained on a confocal microscope with both brightfield and epifluorescence and then overlaid.

### Stability of Plasmid pIGSAF in F. alni ACN14a in the Absence of Selection

In cultures grown without chloramphenicol selection, qPCR analysis did not find a significant difference in the amount of plasmid relative to genomic DNA after one, two, or three rounds of sub-culturing (Figure 5). Only after the fourth round of sub-culturing was a significant decrease from the initial plasmid concentration detected by qPCR (Figure 5), however, fluorescence throughout the hyphae was still observed even after four weeks without selection (Figure 6).

**Figure 5.**
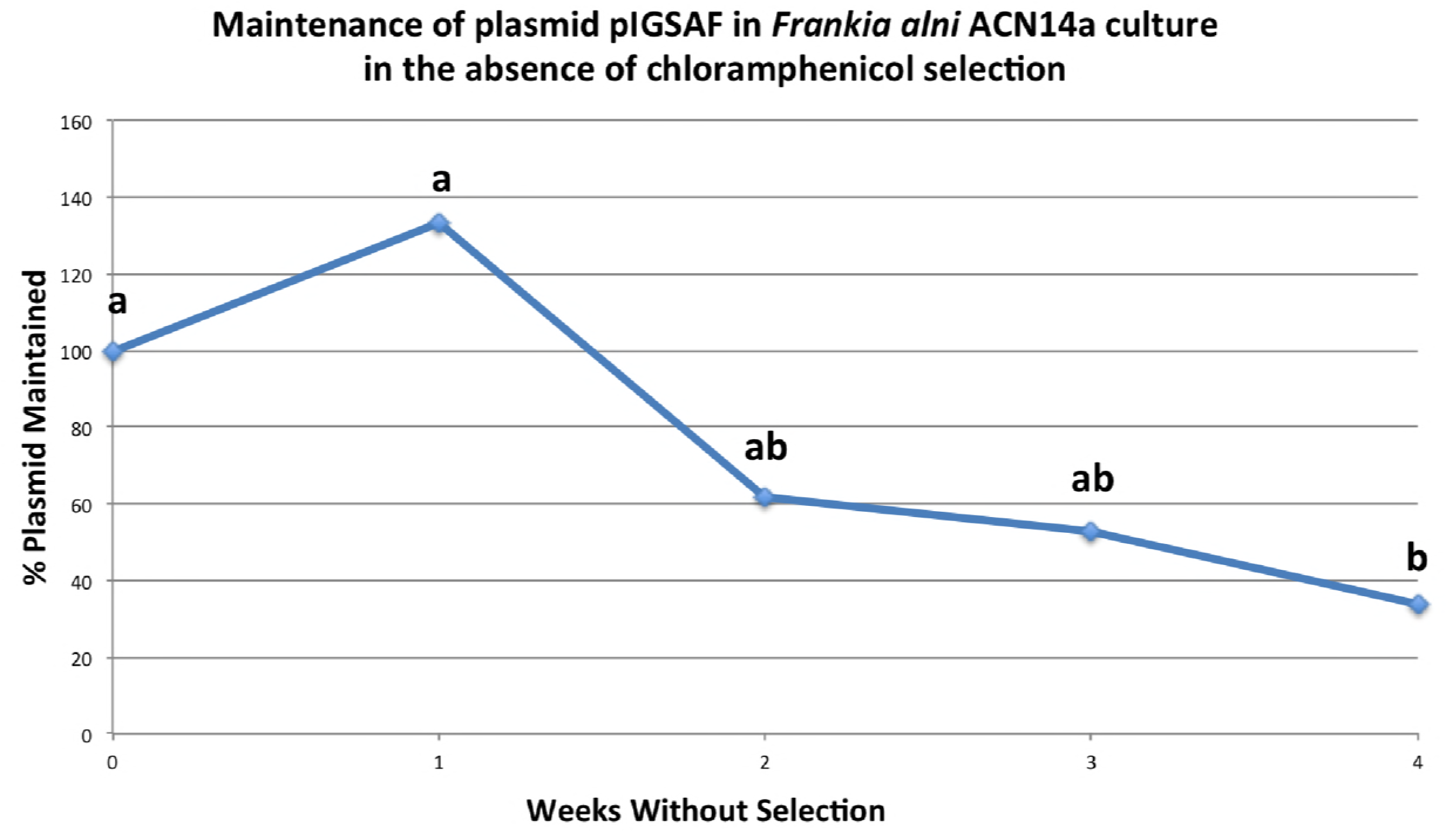
Maintenance of plasmid pIGSAF in *Frankia alni* in culture without chloramphenicol selection over the course of four weeks. Each week the culture was sub-cultured into fresh media and genomic DNA was extracted to measure relative plasmid concentrations via qPCR. Groups ‘a’ and ‘b’ indicate time points that are not significantly different from each other (p<.05).

**Figure 6.**
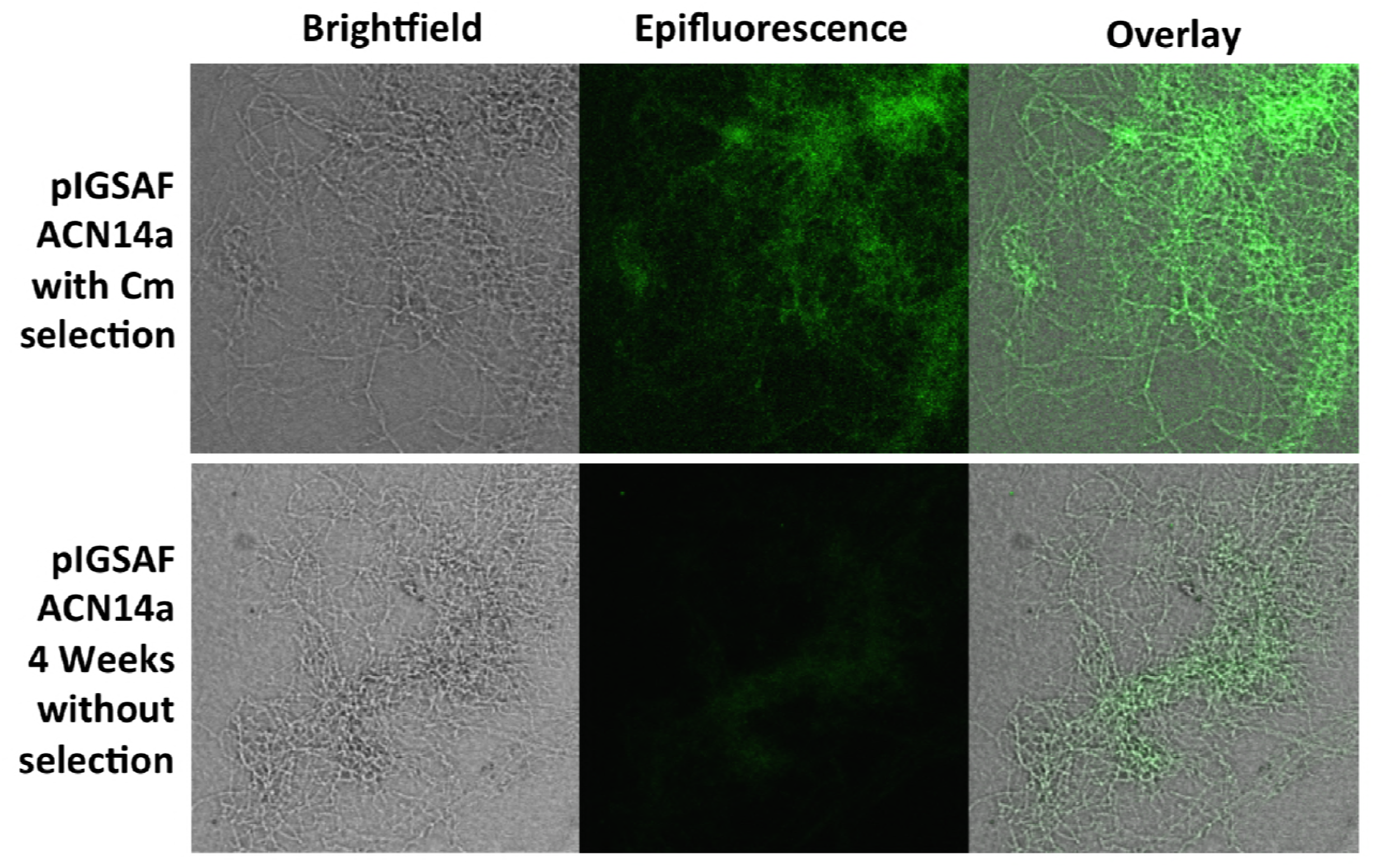
Fluorescence of a *Frankia alni* hyphal colony transformed with plasmid pIGSAF grown with chloramphicol selection (top) and without selection for four weeks (bottom).

## Discussion

### Genetic Transformation of Frankia alni ACN14a with an Unmethylated Replicating Plasmid

We have shown that *F. alni* can be stably transformed with an unmethylated plasmid introduced by electroporation. A homolog of an Mrr Type IV methyladenine-targeting restriction enzyme was highly expressed in *F. alni* in culture (Figure 1), suggesting that DNA with methylated adenine bases would be degraded in this organism. Actinobacteria, especially *Frankia*, expressed Type IV methyl-directed restriction enzyme genes more highly than proteobacteria and firmicutes (Figure 2), a finding that correlates with previous reports of higher transformation efficiencies with unmethylated plasmids in *Corynebacterium* (Ankri *et al*., 1996) and *Streptomyces spp*. (Molle *et al*., 1999). Genomes of the majority of actinobacteria are missing homologs of the *dam* methyltransferase gene (Sachez-Romero *et al*., 2015) whose product is used to mark parent DNA strands during replication, and also are missing *mutS* and *mutL* that form a complex for the removal and repair of mismatched bases on the daughter strand determined by the methylation of adenine residues (Sachadyn, 2010). Together, these factors suggest a difference in preference for unmethylated over methylated DNA among most of the actinobacteria relative to other bacterial phyla.

Type IV restriction enzymes have been suggested to have evolved as a counter to phage methylation systems that themselves evolved to evade host restriction systems by methylating restriction sites (Westra *et al*., 2012). Phage genomes adopt the methylation patterns of their previous host (Loenen and Raleigh, 2014) thus increasing the likelihood of digestion by actinobacterial enzymes if replicated in a *dam*+ host. The expression of type IV restriction enzymes in actinobacteria therefore could represent an adaptation to prevent infection by DNA phages based on the methylation state of their genomes. Differences in methylation patterns between actinobacteria and other bacterial phyla (Novella *et al*., 1996) potentially constitute a barrier to horizontal gene transfer between these groups, including phage-mediated gene transfer. Of particular interest to the evolution of root nodule symbioses is the transfer of genes between *Frankia* and the rhizobia and vice versa. It has been suggested that the *nodA* gene involved in Nod factor synthesis evolved in the actinobacteria, including some *Frankia*, and was then horizontally transferred to the rhizobia (Persson *et al*., 2015). With type IV restriction enzymes creating a barrier to horizontal transfer into actinobacteria from *dam*+ bacteria including proteobacteria, this direction would be more likely than the reverse.

However, *F. alni* was observed to down-regulate its type IV *mrr* gene substantially in symbiosis (Figure 1). As roots contain much lower concentrations of bacteriophage than the surrounding soil (Ward and Mahler, 1982) this likely represents a decreased necessity for restriction enzymes as a defense mechanism during symbiosis. A potential side-effect of this down-regulation, however, is that the barrier to horizontal transfer posed by type IV enzymes is likely lowered during symbiosis, promoting horizontal transfer to *Frankia* from other endophytic bacteria.

*M. tuberculosis* showed much lower transcription of its annotated type IV methyladenine targeting restriction enzyme than other actinobacteria. *M. tuberculosis* expresses an adenine methyltransferase in hypoxic conditions that regulates the expression of genes likely involved with survival during macrophage infection (Shell *et al*., 2013). For this reason it is likely that *M. tuberculosis* responds to methylated DNA differently than other actinobacteria, indeed electrotransformation of *M. tuberculosis* can be readily achieved with methylated plasmids replicated in *E. coli* DH5α (Pelicic *et al*., 1997), suggesting that methylated DNA is not digested in *M. tuberculosis*.

In this study plasmid pSA3 and its derivatives were capable of replication in *F. alni*. This shows that the broad host-range origin is capable of replication in *Frankia* and supports its use as a vector for the manipulation of *Frankia* spp. The parent plasmid of pSA3, pIP501, replicates in a very broad range of bacteria including *Streptomyces lividans* and *E. coli* (Kurenbach *et al*., 2003) indicating the potential for transformation of additional actinobacteria with these plasmids.

### Differential Regulation of egfp Under the Control of the Frankia alni ACN14a nif Cluster Promoter

The expression of the *egfp* gene of plasmid pIGSAFnif was up-regulated in (–)N media compared with expression in (+)N media, at proportional levels similar to the expression of the *nifH* nitrogenase gene (Table 1), demonstrating for the first time that expression of reporter genes can be manipulated in *Frankia*. This transformation system resulted in the ability to visualize the expression of nitrogen-fixation genes *in vitro* by fluorescence microscopy (Figure 4). Interestingly, fluorescence was detected in both the spherical portion of the vesicle as well as in the stalk that connects to the hyphae, suggesting that nitrogen fixation genes are expressed in both parts of the vesicle. Previous studies have shown that the vesicle envelope is deposited around the stalk as well (Lancelle *et al*., 1985), supporting the observation that nitrogen fixation occurs in the stalk.

Although the fluorescence observed when *egfp* was expressed under the control of the *nif* cluster promoter was predominantly in the vesicles, some fluorescence was occasionally observed in hyphae under nitrogen-fixing conditions, whereas in (+)N media there was no observable fluorescence (Figure 4). This suggests that there is some expression of *nif* genes in the hyphae as well as the vesicles induced by nitrogen limitation. *Frankia spp*. in symbiosis with members of the Casuarinaceae have been reported to fix nitrogen in hyphae, since no vesicles are differentiated (Murry *et al*., 1985); this correlated with the formation of a lignified host cell wall in the symbiotic tissue that likely excludes oxygen (Berg and McDowell, 1987). In liquid culture, there may be zones of low pO_2_ that develop in portions of the *Frankia* hyphal colony where nitrogen fixation could be induced.

*Frankia spp*. in symbiosis have been suggested to be more autonomous than rhizobial microsymbionts due to their ability to control the flow of oxygen with the formation of vesicles as well as the expression of more metabolic pathways in the microsymbiont. These factors allow *Frankia* to be more metabolically independent from their hosts (Alloisio *et al*., 2010; Berry *et al*., 2011). As an example, both *Frankia* and the rhizobia share a second glutamine synthetase gene, named *glnII*, which is not present in most other bacteria (Ghoshroy *et al*., 2010). *glnII* is up-regulated in *F. alni* in symbiosis relative to nitrogen-replete culture (Alloisio *et al*., 2010) but not differentially expressed or required for effective nodule formation by rhizobia such as *Sinorhizobium meliloti* (Becker *et al*., 2004; de Bruijn *et al*., 1989). The development of genetic tools for the manipulation of *Frankia* will allow further experimentation into these and other distinctive molecular aspects of actinorhizal symbioses, which will, in turn, inform analyses of the evolution of root nodule symbiosis.

### Future Directions

Even in the absence of selection plasmid pIGSAF was found to be stable in *F. alni* cultures for at least three weeks (Figure 5) and cultures continued to fluoresce after at least four weeks (Figure 6). Three to four weeks is sufficient to nodulate *F. alni* hosts (Alloisio *et al*., 2010), suggesting that transformants can be used to inoculate plants in studies of the role of *Frankia* and its interactions with hosts during nodule establishment and symbiosis. In future this system could be modified using recombination or viral integrases to anchor genes within the genome. Differential regulation of reporter genes such as *egfp* can be used to localize the expression of genes identified by genomics and transcriptomics in specific *Frankia* cell types, in different growth conditions, and in symbiosis. Replicating plasmids may also enable the study of gene function by constitutive expression of selected genomic genes, by promoter switching or by knock-down experiments expressing anti-RNAs to genes of interest (Gillaspie *et al*., 2009). Circumventing the natural restriction systems of *Frankia* will also increase the transformation rate of non-replicating plasmids and enable higher efficiency recombination for gene knock-out experiments as attempted by Kucho *et al.* (2009).

## Materials and Methods

### Restriction Enzyme Analysis

Genes annotated as restriction enzymes (Restriction enzyme types I, II, III, or IV) present in the annotations of all completely sequenced actinobacterial genomes and their annotations were downloaded from the REBASE database (Roberts *et al*., 2010). Bacterial transcriptomes used in this study were downloaded from the NCBI GEO database (Supplementary Table 1). Transcriptomes were chosen based on the following criteria: 1) transcriptomes were made from pure cultures in log-phase growth, 2) organism did not have genetic manipulations including mutations or exogenous plasmids, and 3) cultures did not have additional experimental compounds added including antibiotics or complex carbon sources. For comparisons between transcriptomes, expression levels for each gene were calculated as the percent of genes with lower expression than the gene of interest in each transcriptome.

### Culture Conditions

*E. coli* strains DH5α and GM48 (*dam*- and *dcm*-) were grown in 50ml Difco 1.5% (w/v) Luria Broth (LB) (Catalog #241420, Becton Dickinson, Franklin Lakes, NJ), pH 6.8, in 250mL flasks at 37°C with shaking at 150 rpm overnight. Plates were made with 1.5% (w/v) Bacto Agar (Becton Dickinson, Franklin Lakes, NJ, catalog #214010) in 1.5% LB, and incubated overnight at 37°C. All media were first sterilized by autoclaving for 30 minutes at 121°C.

*Frankia alni* ACN14a (Normand and Lalonde, 1982) was cultured in 5ml liquid BAPP media modified from Murry *et al*., 1984 by the addition of 5mM pyruvate and 5mM MOPS, adjusted to pH 6.7, in sterile 25ml glass test tubes with tetracycline added to a final concentration of 10µg/ml to prevent contamination. (+)N media included 5mM ammonium chloride as a nitrogen source whereas (–)N media had no added nitrogen source. For sub-culturing, hyphae were collected in sterile 2ml microcentrifuge tubes, centrifuged at 10,000rpm for ten minutes in a tabletop microcentrifuge (model #5424, Eppendorf, Hamburg, Germany), and re-suspended in 1mL fresh BAPP media. Cultures were then homogenized by passage through a 21G needle six times, then homogenate equivalent to 50ul packed-cell volume as measured after centrifugation was added to 4ml of fresh media and incubated at 28°C without shaking. Cultures were routinely sub-cultured once per week. For selection of transformants, chloramphenicol was added to a final concentration of 25µg/ml.

### Frankia Transformation

*F. alni* cells were grown in culture for one week prior to transformation. Two milliliters of growth media with hyphae was then pelleted as above and the pellet was re-suspended in 500µL of ice-cold deionized (DI) water that had been sterilized by autoclaving. This was repeated two more times, but on the third round of centrifugation the pellet was re-suspended in 300µL ice-cold sterile 10% glycerol instead. Hyphae in the cell suspension were then homogenized by passage through a 21G needle twice.

300µL of the *F. alni* cell suspension was pipetted into an electroporation cuvette with a 2mm gap (Molecular BioProducts Catalog #5520, San Diego, CA) and mixed with 10µg of plasmid DNA. The cuvette was then incubated on ice for five minutes. Electroporation was carried out in a Bio-Rad Gene Pulser™ with Pulse Controller at 2.5kV, 200Ω resistance, and 25µF of capacitance. The cuvette was then immediately filled with 1ml of ice-cold BAPP media. The cuvette was sealed with Parafilm® (Pechiney Plastic Packaging, Menasha, WI) and incubated overnight without shaking at 28°C.

The following day the *F. alni* culture was removed from the cuvette and added to 3.5ml of sterile BAPP media with tetracycline as above, in a glass test tube. The culture was incubated at 28°C without shaking until visible hyphae were observed (approximately 10 days after electroporation). At this point *F. alni* hyphae were sub-cultured into fresh media as above and then once more one week later. The following week (two weeks after visible hyphae were observed) the hyphae were sub-cultured again into BAPP media, this time with chloramphenicol added for selection to a final concentration of 25µg/ml. Chloramphenicol was chosen as the selective antibiotic because all *Frankia* strains tested in a study by Tisa *et al*. (1998) were susceptible to it. This process was repeated the following week for an additional round of selection.

### DNA Extraction

Plasmids were purified from *E. coli* using a QIAprep® Spin Miniprep Kit (Catalog #27106, Qiagen). Two milliliters of overnight cultures were pelleted at 13,000rpm for five minutes and used for extraction. Plasmids were resuspended in EB buffer (Qiagen) and quantified on a NanoDrop Microvolume Spectrophotometer (ThermoFisher). Plasmids were further purified by running on a 0.7% agarose gel and extracted with a Zymoclean Gel DNA Recovery Kit (Catalog #11-300, Genesee Scientific, San Diego, CA). To synthesize unmethylated plasmids, methylated plasmids were first extracted from *E. coli* DH5α and transformed into *E. coli* GM48 by heat shock in CaCl_2_ (Sambrook *et al.*, 1989) then cultured and re-extracted as above.

*F. alni* DNA extraction was carried out with a CTAB protocol (Feil *et al*., 2012). Briefly, cells were pelleted at 10,000rpm for five minutes in a tabletop centrifuge. Cells were then re-suspended in TE buffer and lysed with lysozyme, SDS, Proteinase K, and CTAB. DNA was extracted with 24:1 chloroform:isoamyl alcohol, and 25:24:1 phenol:chloroform:isoamyl alcohol, washed with isopropanol followed by ethanol, resuspended in 50ul TE buffer, and quantified by NanoDrop.

### Plasmid Synthesis

A plasmid designated as pIGSAF was synthesized by ligating a PCR-amplified fragment containing the *egfp* gene and promoter from plasmid pDiGc (Helaine *et al*., 2010, Addgene, Cambridge, MA) to plasmid pSA3 (Dao and Ferretti, 1985). Plasmid pSA3 is a broad host-range replicating plasmid developed as a shuttle vector from pIP501. The pIP501 origin of replication has been shown to replicate in bacteria of diverse phyla including Firmicutes, from which it was originally isolated (Horodniceanu *et al*., 1976), as well as Actinobacteria and Proteobacteria (Kurenbach *et al*., 2003). Plasmid pSA3 contains chloramphenicol and tetracycline resistance genes and an *E. coli*-specific origin of replication for propagation (Dao and Ferretti, 1985).

Primers for PCR, listed in Table 2, were designed using NCBI Primer-BLAST with default settings (Ye *et al*., 2012). For cloning, primers were designed with linkers adding target restriction sites onto their 5’ ends (Table 2) as well as 4-6 additional bases to aid in restriction digestion of the ends. A diagram of the addition of linkers by PCR is shown in Figure 7A. For the synthesis of pIGSAF, primers were designed targeting the *egfp* coding region as well as the promoter region 200 base pairs upstream. PCR was performed in a Bio-Rad S1000 Thermal Cycler (Bio-Rad, Hercules, CA) using a Qiagen Taq PCR kit (catalog #201223). Products were synthesized as in Figure 7A, first amplified for 10 cycles using the annealing temperature of the region corresponding to the binding site on the target DNA and then an additional 30 cycles using the annealing temperature of the full primer including the linker (Table 2).

**Table 2.**
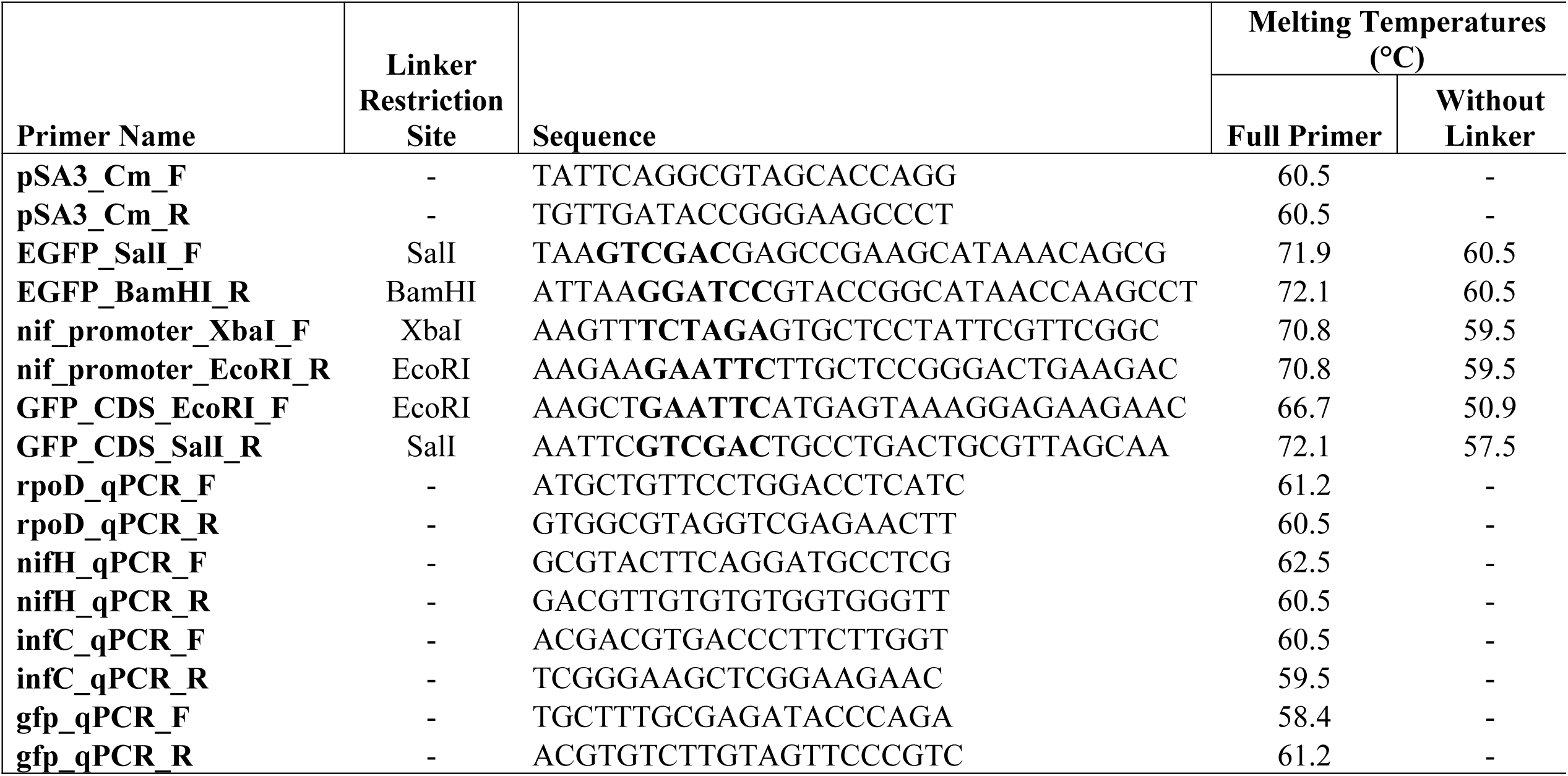
Primers designed for this study. Restriction sites added to primers by linker PCR for use in cloning are bolded in the primer sequence and annealing temperatures are listed both for the initial reaction (“Without Linker”) and the main amplification phase of the PCR (“Full Sequence”).

**Figure 7.**
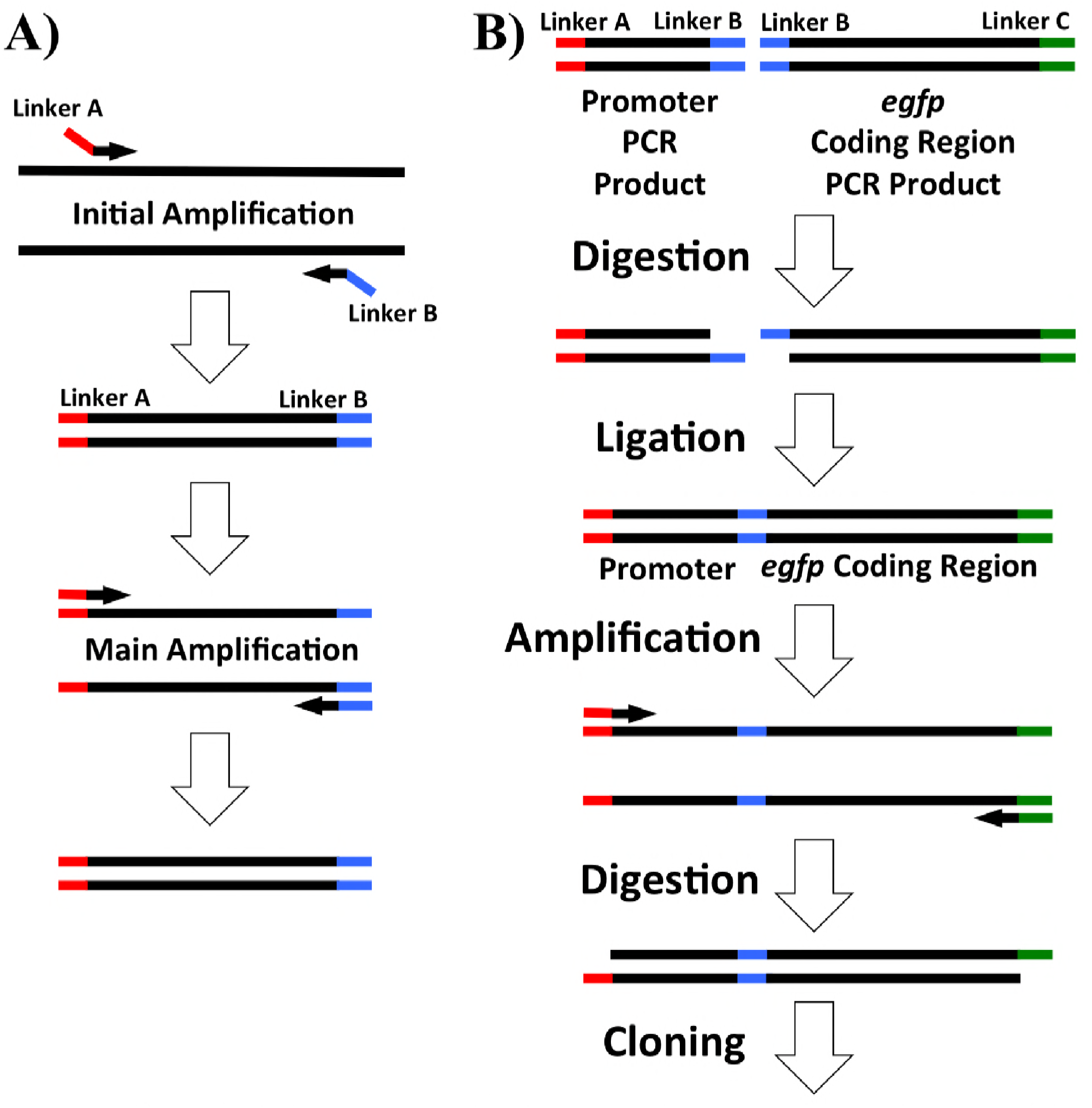
PCR amplification and digestion and ligation overview for plasmid synthesis used in this study. Linkers including restriction sites are colored red, blue, or green. A) PCR products for cloning were amplified by amplification at a lower initial temperature to allow the incorporation of additional restriction sites on the 5’ end of the primers (shown in red and blue). The annealing temperature was then increased to amplify full-length products with the added restriction sites. B) Addition of the *F. alni* ACN14a *nif* cluster promoter region to the *egfp* coding sequence. Both the promoter region and *egfp* coding sequence were amplified with EcoRI sites (blue). These were then digested and ligated together followed by amplification of the ligation product. Restriction sites on the ends (red and green) were digested, allowing incorporation of the *nif* promoter:*egfp* product into the digested plasmid.

The *egfp* fragment was synthesized with SalI and BamHI restriction sites on the 5’ and 3’ ends, respectively, with primers EGFP_SalI_F and EGFP_BamHI_R. The PCR product and plasmid pSA3 were then digested with both enzymes. The fragments were purified on an agarose gel as above and then mixed together, denatured by heating for five minutes at 65°C, and then ligated together by incubation with T4 ligase (catalog #M0202S, Qiagen) at 16°C overnight. The resulting ligation was transformed into *E. coli* DH5α by heat shock (Sambrook *et al.*, 1989). Transformants were selected on LB plates with chloramphenicol at a final concentration of 25µg/ml. Transformed colonies were inoculated into liquid LB media with chloramphenicol and cultured as above. Plasmid pIGSAF was then re-extracted from transformed *E. coli* and its composition was confirmed by digestion with SalI and BamHI as above.

The presence of plasmids pSA3 and pIGSAF electroporated into *F. alni* cultures was confirmed with PCR. Genomic DNA was extracted and amplified with primers pairs pSA3_Cm_F/pSA3_CmR for the chloramphenicol resistance gene of plasmid pSA3 and GFP_qPCR_F/GFP_qPCR_R for the *egfp* gene of plasmid pIGSAF. PCR products were separated on an agarose gel and extracted as above, and sequenced by the UCDNA Sequencing Facility (Davis, CA). Sequences were compared by BLAST against gene sequences obtained from the original plasmids (Supplementary Table 2).

For the differential expression of *egfp* by nitrogen limitation under the control of the *nif* cluster promoter region of *F. alni* ACN14a, plasmid pIGSAFnif was synthesized. Ligation of the *nif* cluster promoter region to the coding region of *egfp* was carried out as outlined in Figure 7B. The *egfp* coding region of pDiGc was amplified without the upstream promoter region using primers GFP_CDS_EcoRI_F and GFP_CDS_SalI_R (Table 2). An EcoRI restriction site was added before the start codon and a SalI site was added 200 bases downstream of the stop codon. Separately, the 300 bases upstream of the *nif* nitrogenase cluster in the *F. alni* ACN14a genome were amplified with an XbaI site upstream and an EcoRI site downstream using primers nif_promoter_XbaI_F and nif_promoter_EcoRI_R (Table 2). These two PCR products were digested with EcoRI (catalog #R0101S, New England Biolabs) and ligated together. The ligation product was then re-amplified by PCR using the *nif* promoter forward primer and the *egfp* reverse primer. The amplified ligation product and plasmid pSA3 were then each digested with SalI and XbaI, ligated together, transformed, and selected on chloramphenicol as above.

### qPCR Verification of Differential Regulation

Quantifications of gene expression and fold-changes were obtained based on qPCR amplification. Cultures of transformed cells were grown in 4.0 ml (+)N and (–)N media in sterile six-well plates (catalog #353046, Corning Inc, Corning, NY) with shaking at 50rpm at 28°C. After five days, RNA was extracted by bead-beating, following a protocol adapted from Dietrich *et al*. (2000): *F. alni* hyphae were pelleted at 9000rpm for fifteen minutes, resuspended in 1050ul Buffer RLT (Qiagen), and transferred to 2ml tubes containing Lysing Matrix B (catalog #6911-100, MP Biomedicals, Burlingame, CA). Samples were processed with a FastPrep FP120 (Thermo Fisher Scientific, Waltham, MA) for 45 seconds at setting 6.5, then placed on ice for 45 seconds. The processing step was then repeated twice more with cooling on ice between each step. Supernatants were then transferred to Qiagen RNeasy spin columns and purified with a Qiagen RNeasy Mini Kit (catalog #74104). RNA was eluted in RNase-free water and then contaminating DNA was digested with an Invitrogen TURBO DNA-free Kit (catalog #AM1907, Waltham, MA). Finally, cDNA was synthesized with an Invitrogen Superscript III Kit (catalog #18080051) with random hexamer primers. qPCR was performed on a 7500 Fast Real-Time PCR System (Applied Biosystems, Foster City, CA) with Fast SYBR Green qPCR Master Mix (catalog #4385612, Applied Biosystems) and primer pairs rpoD_qPCR_F/rpoD_qPCR_R, nifH_qpCR_F/nifH_qpCR_R, infC_qPCR_F/infC_qPCR_R, and gfp_qPCR_F/gfp_qPCR_R (Table 1). *egfp* and *nifH* were used as experimental targets to confirm differential regulation of *egfp* by nitrogen limitation; housekeeping gene *infC* was used for normalization as in Alloisio *et al*. (2010); and sigma factor gene *rpoD* was used as a negative control.

### Confocal Microscopy

For visualization of hyphae and GFP fluorescence, *Frankia* cells were grown either in (+)N or (–)N BAPP medium as above and immobilized on glass slides with a drop of 3% molten agarose solution: 1.5g molecular biology-grade agarose (catalog #A9539, Sigma, St. Louis, MO) was dissolved in 50ml DI H_2_O and kept warm in a water bath at 50°C. Slides were pre-heated on a slide warmer (Fisher) at 50°C. Then 15ul of *Frankia* hyphae were pipetted onto each slide and covered with 35ul of 3% molten agarose. A #1.5 coverslip was added to the *Frankia* cells in agarose, which were allowed to cool to room temperature. The *Frankia* preparations were visualized on a Leica TCS SP8 STED 3X confocal microscope with a 100X oil-immersion objective and a HyD detector. For fluorescence imaging, samples were excited with 488nm light. The emission wavelengths were collected from 500-550nm. Images were stored as. lif files from Leica LAS X and then viewed in FIJI (Schindelin *et al*., 2012). To visualize hyphal three-dimensional structure, Z-stack images were taken and then combined using the highest fluorescent intensity of each pixel (FIJI MAX setting).

### Plasmid Stability

In order to test the persistence of a plasmid in transformed *F. alni*, cultures containing plasmid pIGSAF were grown without selection in non-selective media lacking chloramphenicol. The cells were first pelleted and re-suspended in fresh BAPP media, then sub-cultured as above without the addition of chloramphenicol to the media. These cultures were grown in six-well plates as above for one week. At that time each culture was pelleted and re-suspended in 500ul fresh BAPP media. This suspension was homogenized by passage through a 21G needle twice. 250ul of the homogenate was transferred to fresh BAPP media without chloramphenicol and the remaining 250ul was used for total genomic DNA extraction as above. This process was repeated once per week for four weeks. The amount of plasmid in each sample was quantified by qPCR, performed in duplicate for each of three biological replicates, using *egfp* primers GFP_qPCR_F and GFP_qPCR_R (Table 2). Relative fold-change of plasmid between each time point was calculated using the ΔΔCt method (Lee *et al*., 2006) with the *infC* gene as a control to normalize the amount of DNA in each sample (amplified with primers infC_qPCR_F and infC_qPCR_R, Table 1). To determine significant changes in plasmid abundance, two-tailed Welch’s t-tests were performed in R on normalized ΔCt values calculated (p<.05). Cultures grown with selection and without selection for four weeks were also imaged with fluorescence as above.

## Acknowledgements

We thank Dr. Rebecca Parales for providing plasmid pSA3 and Dr. Wolf Heyer for *E. coli* strain GM48. We also thank Dr. Ingrid Brust-Mascher for technical assistance with confocal microscopy. IG was supported by a UC Davis Department of Plant Sciences graduate student research fellowship. Research funding for this project was provided by UC Davis Henry A. Jastro Research Grants to IG, and by USDA-NIFA-CA-D-PLS-2173-H (AB).

## Supplementary Figures and Tables

**Supplementary Table 1** List of genomes and transcriptomes used in this study. Transcriptomes are listed with their GEO accession number and original reference.

**Supplementary Table 2** Sequences from PCR products of *Frankia alni* transformants corresponding to the chloramphenicol resistance gene (*camr*) from pSA3 and *egfp* gene of plasmid plasmid pIGSAF and pDiGc.

